# BIND: Large-Scale Biological Interaction Network Discovery through Knowledge Graph-Driven Machine Learning

**DOI:** 10.1101/2025.01.15.633109

**Authors:** Naafey Aamer, Muhammad Nabeel Asim, Aamer Iqbal Bhatti, Andreas Dengel

## Abstract

The complex interactions between biological entities provide valuable insights into fundamental life processes, which pave the way to a deeper understanding of disease mechanisms for the development of innovative therapeutic strategies. To enable large-scale predictions of biological interactions, multifarious AI-driven predictors have been developed. However, most of these are developed by leveraging information from only a limited subset of interaction types, and the broader interaction types landscape could facilitate AI algorithms to learn more informative patterns to predict unknown interactions. To address the need of a robust and precise interaction predictor, we introduce BIND (Biological Interaction Network Discovery), a predictor that leverages simultaneous learning across 10 biological entities and 30 interaction types. To develop BIND, we first evaluate 11 distinct Knowledge Graph Embedding Methods on the largest public biomedical interaction dataset namely PrimeKG. For each relation type, we extracted entity embeddings from the top 5 performing Knowledge Graph Embedding Models (KGEMs) and input them into 7 distinct machine learning classifiers. Rigorous evaluation of 1,050 predictive pipelines demonstrated that specific combinations of KGEMs and classifiers achieved F1 scores of 90% to 99% across various interaction types. Comprehensive evaluation across each relation type identified the top-performing predictive pipelines, which became the foundation of the BIND web application. To reveal the practical utility of our web application in identifying novel biological interactions, we conducted a case study on drug-phenotype interactions. The application gave 1,355 high confidence predictions, from which potential interactions were subsequently validated by scientific evidence found within the existing literature (Table 5). We believe BIND’s web application’s public access will serve as a valuable tool for biologists to identify unknown interactions that can be subsequently validated through wet-lab experiments.

## Introduction

Biological systems are derived from complex interactions between entities at variable scales ranging from small biomolecules like genes and proteins to macroscopic structures like organs and tissues. A profound understanding of these complex hierarchies and interactions is essential for developing innovative solutions to address human health challenges. Such biological interaction data naturally takes the form of a network/graph, where each type of interaction offers unique insights into biological processes and provides potential avenues for therapeutic interventions in biomedicine. Notably, protein-protein interactions reveal critical cellular pathways and potential drug targets (1) (2); disease-drug interactions illuminate therapeutic responses and opportunities for drug repositioning (1) (2); disease-gene interactions uncover diagnostic biomarkers and genetic predispositions that guide treatment approaches for complex diseases like Alzheimer’s and Autism (3) (4); drug-drug interactions provide crucial insights into treatment synergies, adverse effects, and optimal therapeutic combinations (2) (1); and disease-disease interactions highlight shared pathological mechanisms and comorbidity patterns (1, 5). Collectively, these diverse interactions form an intricate web of biological relationships that, when analyzed comprehensively, enables the elucidation of complex disease mechanisms, and accelerates the discovery of novel biomarkers(2). While traditionally these interactions are identified through wet-lab experiments, such approaches are expensive, time-consuming, and error-prone (6, 7). This has led to a paradigm shift towards computational predictors that harness the potential of Artificial Intelligence (AI) for interaction prediction (8). Moreover, knowledge graph embedding methods (KGEMs) have enabled the development of efficient and large scale interaction prediction applications.

These AI predictors make use of a graphical space where molecules, drugs, genes, proteins, and diseases are represented as nodes and the interactions between them are denoted through edges (9–11). This space is commonly derived from biological knowledge graphs. A mainstream method for utilizing knowledge graphs is to learn low dimensional feature vectors where structurally similar nodes have similar encoded vectors and dissimilar nodes have distinctly encoded vectors (10, 12, 13). These encoded vectors (embeddings) are utilized to train AI classifiers for downstream interaction prediction tasks. These knowledge graph embeddings-based AI predictors can effectively solve complex problems where both relationship structures and individual molecular properties are important (9, 14).

However the full potential of the knowledge graph embedding methods (KGEMs) has not been explored as current research typically tends to operate in isolation, focusing on single tasks like drug-target interactions (15)(16), disease associations (17)(18), drug repurposing (19)(20), and protein interaction prediction (21–23). These approaches may miss the broader picture of how different biological interactions influence each other. For example, protein-protein interactions can shape drug-disease relationships, missing proteins can impact anatomical structures, and pathway interactions can reveal new drug repurposing opportunities. To capture these complex interactions, one could leverage large-scale biological interaction datasets which have a large number of relation types. However, this approach introduces significant noise that could hurt prediction reliability, and traditional solutions like extensive ablation studies are computationally prohibitive for such large datasets. Our solution lies in adopting a strategic finetuning approach: first training the AI predictor globally on the entire dataset to capture the full context, then carefully finetuning it on the specific relation subset of interest. This approach allows us to leverage the full context while maintaining focused predictions. While the need for a comprehensive solution is clear, no unified platform currently exists where biologists can predict and analyze multiple types of biological relationships together. This gap prevents researchers from fully understanding how different biological mechanisms interact and limits our ability to discover new therapeutic applications.

The primary objective of this research is to address these limitations with a unified platform with state-of-the-art predictive pipelines that can accurately predict distinct types of interactions between different biological entities. To explore and achieve this objective, we propose BIND, **B**iological **I**nteraction **N**etwork **D**iscovery, a comprehensive framework and web application for unified biological interaction prediction. BIND is developed on the basis of large scale experimentation to find the optimal training environment for 1050 unified predictive pipelines based on 11 Knowledge Graph Embedding methods and 7 Machine Learning classifiers that are trained and evaluated across 30 distinct types relations. The top-performing embedding method + classifier combinations for each relation are deployed on our web application https://sds-genetic-interaction-analysis.opendfki.de/. Originating from extensive experimentation involving several hundred experiments with 1,000+ GPU hours and 15,000+ CPU hours of computation time, a summary of our contributions are summarised as follows:

1. A comprehensive evaluation of 11 Knowledge Graph Embedding Methods (KGEMs) on 8 million interactions between 30 biological relationships and 129 thousand nodes, revealing that architecturally simpler models effectively capture the inherent nature of biological interactions, often outperforming more complex approaches.
2. Effective representation learning with a two-stage training strategy where:
  - Initial training on all 30 interaction types simultaneously captures complex inter-relationships between different biological interactions
  - Relation-specific fine-tuning is done to optimize the embeddings from the first stage for each interaction type while preserving broader biological context, achieving improvements up to 26.9% for protein-protein interactions.
3. Identification of optimal embedding-classifier combinations for each biological interaction type, resulting in optimal predictive performance (F1-scores: 0.90-0.99) across different biological domains, with specific combinations showing superior performance for different relation types.
4. BIND (Biological Interaction Network Discovery), a unified web application that:
  - Searches for novel interactions from billions of potential relationships across 30 biological interaction types with optimal embedding-classifier combinations for each relation type
  - Deploys these optimized models in a comprehensive web application, providing a one-stop platform for biological interaction prediction and discovery
  - Demonstrates practical utility through a case study that successfully validated novel drug-phenotype interactions from literature out of the 1,355 high confidence predictions, offering a publicly accessible tool for biologists to identify unknown interactions for experimental validation.

## Materials And Methods

This section presents the core components of BIND’s methodology. It first describes PrimeKG, the benchmark dataset beneath our framework, followed by a discussion of Knowledge Graph Embedding Methods (KGEMs), with particular emphasis on decomposable architectures and their applicability to large-scale biomedical knowledge graphs. Next, our choices for machine learning classifiers are discussed and the section concludes with the evaluation metrics used to assess both the embedding quality and classification performance of the framework.

### Benchmark Dataset

The biological interaction prediction landscape has witnessed the development of multifarious knowledge graph datasets including SPOKE(24), HSDN(25), GARD(26), PrimeKG(27), and BioKG(28). Among these datasets, PrimeKG has several key advantages that make it particularly well-suited for the development of AI-driven interaction prediction applications. With 22,236 disease terms consolidated into 17,080 clinically meaningful conditions, it surpasses the count of diseases in previous datasets by 1-2 orders of magnitude(27). Furthermore, PrimeKG contains a broad range of 30 distinct relation types that have been designed to specifically cater disease and drug information with specialized relations like indication, contraindication, and off-label use(27). PrimeKG’s explicitly defined bidirectional relations and coverage across all primary biomedical subdomains enable learning representations that capture disease interactions in a truly global context. PRIMEKG represents a significant advancement in the development of AI-powered precision medicine applications by integrating data from 20 diverse sources, including DrugBank(29), DisGeNET(29), CTD (Comparative Toxicogenomics Database)(30), Gene Ontology(31), Human Phenotype Ontology(32), and Reactome(33). These advantages led us to select PrimeKG as the base of BIND.

The dataset comprises 129,375 nodes across 10 types, including drugs, diseases, genes/proteins, biological processes, molecular functions, cellular components, anatomical structures, pathways, effects/phenotypes, and environmental exposures. The graph contains 4,050,249 unique relationships (8,100,498 total when accounting for bidirectionality) with explicitly defined bidirectional relations, requiring models to capture both forward and reverse relationships effectively.

Although PrimeKG is a large-scale dataset, it presents multiple challenges for predictor development. Most notably, it exhibits significant skewness in its relation distribution - two relation types, Anatomy-Protein (37.5%) and Drug-Drug interactions (33%), account for over 70% of all relations, while critical biological relationships like Disease-Phenotype (3.7%) and Protein-Protein interactions (7.9%) remain sparse. A more detailed information about relations imbalance can be seen in Figure 2. This stark imbalance requires efficient models capable of handling both dense and sparse relationships effectively. PrimeKG’s architectural heterogeneity adds another layer of complexity, with some relations forming dense, interconnected subgraphs (like Drug-Drug interactions) while others exhibit hierarchical structures (like Disease-Phenotype). The structural diversity extends to relation cardinalities, ranging from one-to-one mappings in drug interactions to many-to-many relationships in Anatomy-Protein associations. Furthermore, PrimeKG’s comprehensive coverage spans both rare diseases (90.8% of Orphanet’s 9,348 rare diseases) and common conditions, requiring models to effectively handle both frequent and rare entity occurrences.

**Fig. 1.**
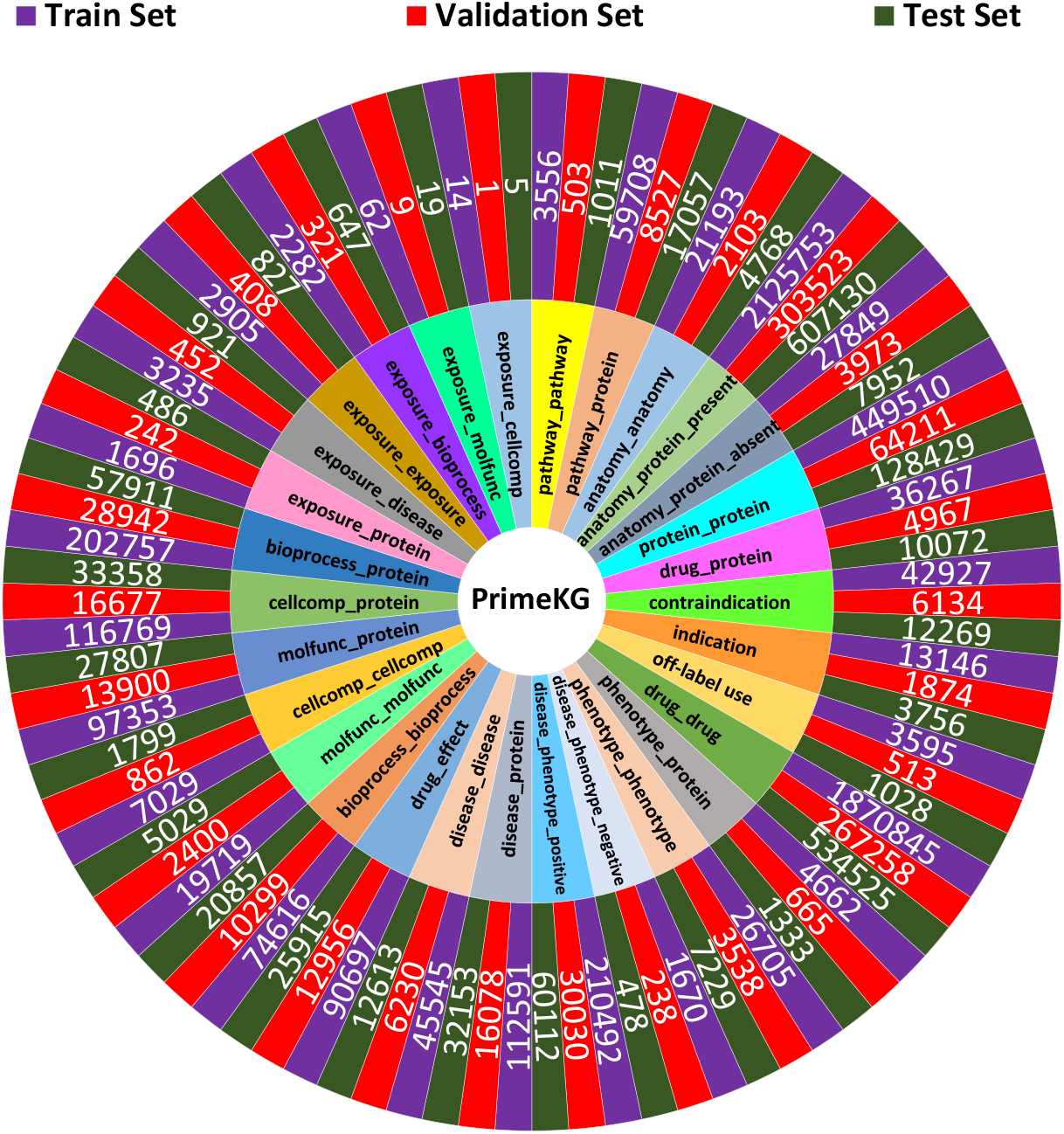
The distribution of PrimeKG’s 30 relations in train, test and validation sets.

**Fig. 2.**
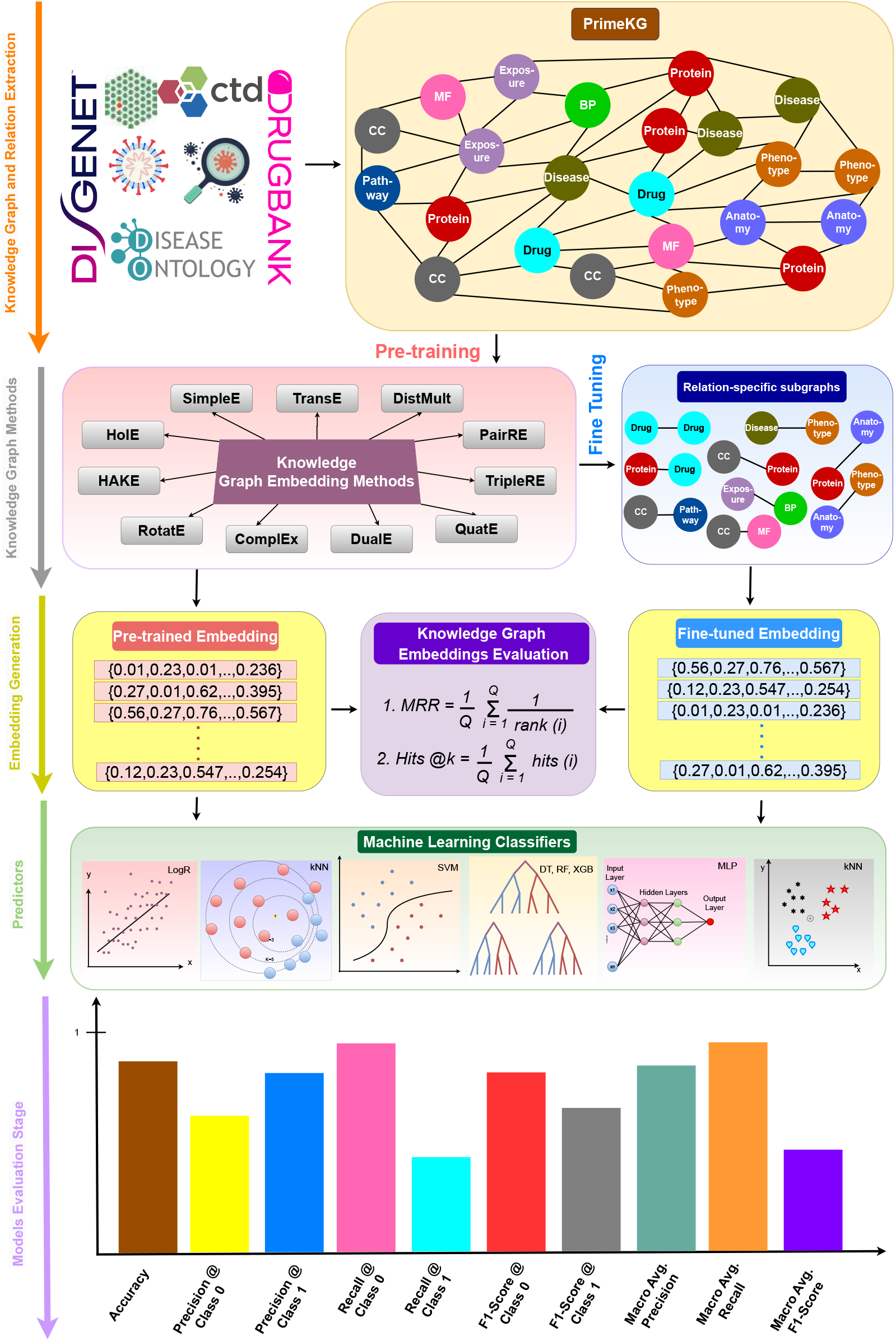
A high-level representation of the distinct modules of the BIND framework

To facilitate model development and evaluation, we partition PrimeKG into training, validation, and test sets following a 70:10:20 ratio, resulting in 5,675,168 training triples, 807,834 validation triples, and 1,617,496 test triples. These splits are stratified for each relation separately to ensure equal distribution of relations in each set. The relation wise splits are shown in Figure 2. The training set is used for model optimization, while the validation set guides hyperparameter selection and early stopping decisions. The test set is reserved exclusively for final model evaluation and is only accessed after completing all model development and tuning phases.

### Knowledge Graph Embedding Methods

Knowledge Graph Embedding Methods (KGEMs) can be broadly categorized into two architectural paradigms: monolithic and decomposable models (45). Monolithic models, such as ConvKB (46) and KBGAT (47), employ deep neural architectures that allow for arbitrary interactions between entities and relations, taking the form:

While these models offer greater flexibility in capturing complex relationships and have demonstrated better performance on smaller benchmark datasets like FB15K(34) and WN18RR(48), they face significant challenges for largescale biomedical knowledge graphs. Their computational requirements and memory footprint make them prohibitively expensive to train on datasets like PrimeKG, which is typically an order of magnitude larger than traditional benchmarks. Moreover, their architectural complexity makes it difficult to conduct comprehensive hyperparameter exploration and ablation studies necessary for understanding model behavior in biomedical contexts. These inherent challenges make it difficult to develop robust applications in the biological interaction landscape.

In contrast, decomposable models learn informative patterns from large knowledge graphs with more ease and provides more opportunities to predict unknown relations. They offer a more tractable approach while maintaining competitive performance (49). These models restrict interactions to elementwise operations between relation-specific subject and object embeddings, taking the general form:

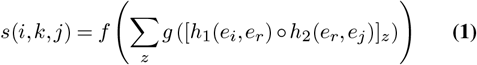

In Equation 1, *∘*represents element-wise operations, *h*_1_ and *h*_2_ transform entities based on the relation, and *g* and *f* are scalar functions. Consider DistMult (35), one of the most fundamental decomposable models. DistMult represents each entity *i* as a vector **e**_*i*_∈ℝ^*d*^ and each relation *k* as a diagonal matrix **R**_*k*_ ∈ℝ^*d×d*^. For a triple (*i, k, j*), it computes the score as:

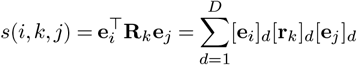

This scoring mechanism can be interpreted as measuring the similarity between the head entity and tail entity in a relation-specific space, where the relation determines the importance of each dimension through its diagonal matrix. The simplicity of this approach allows for efficient computation and straightforward interpretation, while still capturing meaningful patterns in the data.

Most decomposable models follow this general paradigm, primarily differing in their specific scoring functions and the geometric intuitions they encode. For instance, TransE(34) interprets relations as translations in the embedding space, while RotatE(41) represents them as rotations in complex space. Despite their architectural constraints, these models have demonstrated remarkable effectiveness, often matching or exceeding the performance of more complex architectures when trained with modern optimization techniques (50)(51). Given these considerations, our work focuses exclusively on decomposable models. This choice is motivated by three key factors: (1) their computational efficiency enables comprehensive experimentation on large-scale biomedical knowledge graphs, (2) their interpretable nature aligns well with the requirements of biomedical applications where model decisions often need to be explained - this interpretability stems from their reliance on simple similarity measures for learning, making them significantly more transparent than black box deep neural networks - and (3) their competitive performance suggests that the additional complexity of monolithic models may not be justified for our specific use case.

Table 1 shows the decomposable Knowledge Graph Embedding Models (KGEMs) that are evaluated for BIND. The model selection is guided by their proven influence in biomedical applications (52)(53). While we prioritized established models that introduced significant novelties, we also included recent architectures to ensure a fair and comprehensive evaluation.

**Table 1.**
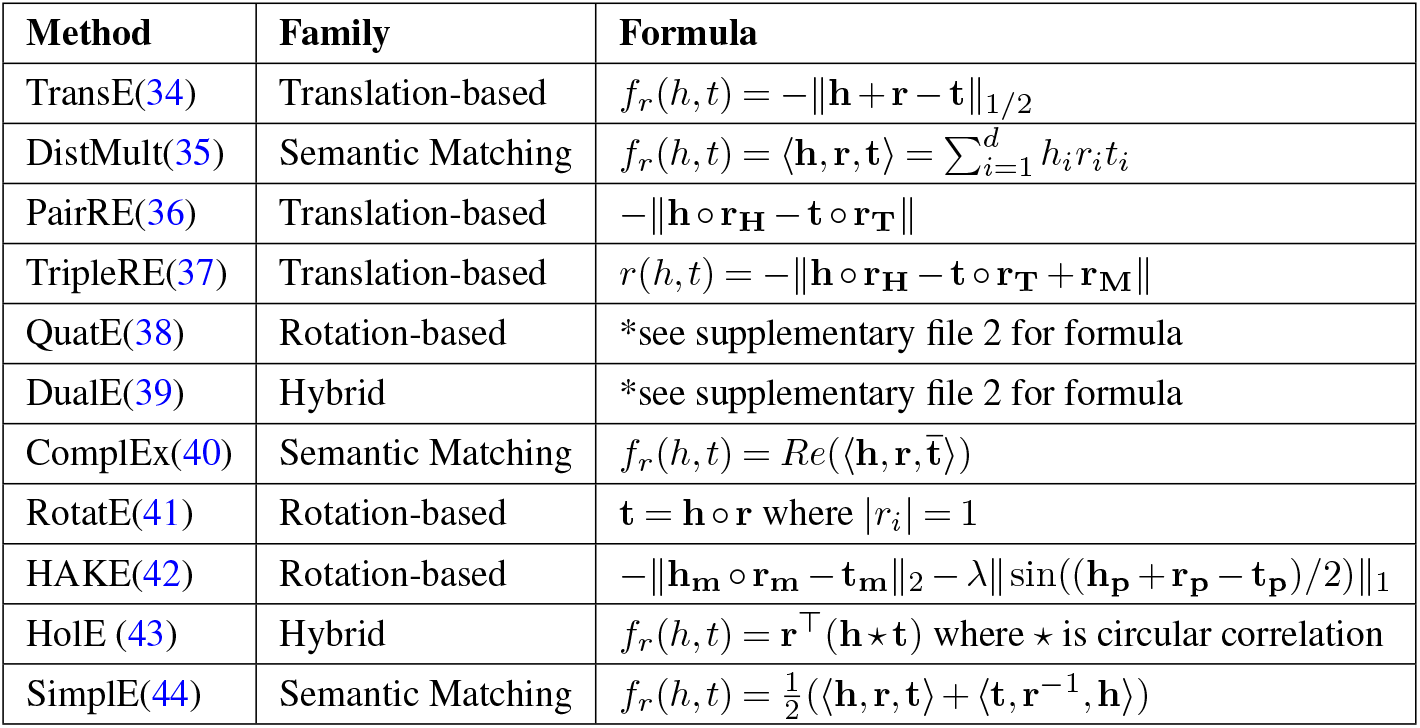
Knowledge Graph Embedding Methods and Their Scoring Functions.

These methods can be categorized into 4 distinct families: translation-based models (TransE(34), PairRE(36), TripleRE(37)), rotation-based models (RotatE(41), QuatE(38)), semantic matching models (DistMult(35), ComplEx(40), SimplE(44), HAKE(42)), and hybrid approaches (DualE(39), HolE(43)). TransE pioneered the translational approach where relationships are interpreted as translations in the embedding space, while PairRE and TripleRE extended this with pair-wise and triple-wise relation embeddings respectively. The rotation family, including RotatE and QuatE, models relationships as rotations in complex or quaternion space, with HAKE(42) further incorporating hierarchical structure through modulus information. The semantic matching models (DistMult, ComplEx, SimplE) use various multiplication-based scoring functions, where ComplEx extends DistMult to complex space for better asymmetric relation modeling. DualE(39) and HolE(43) represent hybrid approaches, with DualE combining rotation and translation, while HolE employs circular correlation for capturing compositional relationships.

### Classifiers

The BIND framework’s classification module encompasses 7 machine learning classifiers, carefully selected to provide a comprehensive evaluation of different prediction approaches for each type of biological interaction. These classifiers span the fundamental paradigms of machine learning: linear classifiers like Logistic Regression(54) operate on direct feature combinations; distance-based methods such as K-Neighbors Classifier(55) and SVM(56) capture spatial relationships in the embedding space; tree-based ensembles including both parallel (Random Forests(57)) and sequential boosting approaches (XGBoost(58)) leverage hierarchical decision boundaries; and neural network classifiers through MLP(59) learn complex feature interactions. The Decision Tree(60) serves both as a standalone method and a building block for ensemble approaches. This set of classifiers allows us to thoroughly analyze how different learning algorithms perform across the 30 distinct types of biomedical relations when combined with various knowledge graph embedding methods.

### Evaluation Measures

The best performing embedding methods and classifiers are chosen for the development of BIND by a comprehensive evaluation criteria encompassing both knowledge graph embeddings (KGEs) and downstream classification tasks.

Following evaluation criteria of existing KGEs studies(61), we utilized two primary metrics: Mean Reciprocal Rank (MRR) and Hits@K, where K = {1,3,5,10,50}. MRR quantifies the average reciprocal rank of correct entities across test set triplets, while Hits@K measures the proportion of test set triplets where the correct entity appears in the top K predictions. These metrics evaluate the model’s ability to rank correct entities in the embedding space, as formalized in Equation 2.

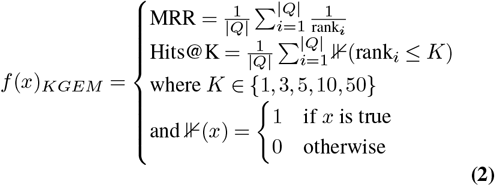

For classification tasks, we employed standard, micro, and macro versions of four key metrics: accuracy (Acc), precision (Pr), recall (R), and the F1-score (F1). The standard formulations are presented in Equation 3.

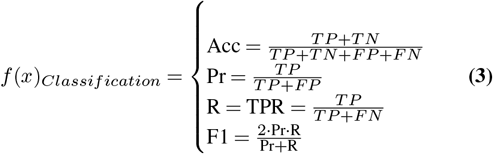

### Experimental Setup

The BIND framework is implemented by leveraging Python 3’s multiprocessing capabilities. For Knowledge Graph Embedding Methods (KGEMs), implementations are acquired either from authors’ original codebases or through the Py-Keen library (62). These implementations are subsequently optimized and parallelized for GPU execution using PyTorch 2.0 (63). For downstream classification tasks, Intel-optimized Scikit-learn (64) is utilized, which offers highly efficient CPU parallelization for classical machine learning algorithms. The web application is developed using Django with PostgreSQL. The details about the underlying dataset and its partitioning can be found in Section 2.

### Training Methodology

BIND’s framework combines a pool of knowledge graph embedding methods and Machine Learning classifiers to create and evaluate end-to-end predictive pipelines through three distinct training phases.

In the first phase, 11 KGEMs are trained on the complete train set encompassing all 30 interaction types. The training procedure implements early stopping with a patience of 5 epochs, monitored on the validation set’s Mean Reciprocal Rank (MRR). A GPU-friendly training process is utilized that employs chunking for memory conservation, automatic mixed precision for efficient computation, and gradient clipping to ensure stable optimization.

The second stage involves the implementation of a targeted finetuning strategy to optimize performance for individual relations. For each relation, the top 5 performing models are identified based on test set MRR, which are then used for relation-specific training. This process involves loading the pre-trained models and further optimizing them exclusively on the subset of training triples that contain the target relation. This finetuning phase yields embeddings from 150 distinct models (30 relations × 5 KGEMs per relation).

These embeddings serve as input features to 7 distinct Machine Learning classifiers which perform binary interaction prediction. For this, a filtered negative sample is generated that corresponds to every positive edge in the training set, ensuring balanced class distribution. The final phase evaluates interaction prediction using seven distinct classifiers, resulting in 1,050 unique predictive pipelines (30 relations × 5 KGEMs × 7 classifiers).

### Loss

For the first two phases, the training objective is optimized using Cross-Entropy Loss (CEL) computed over the entire training set. For each triple (*h, r, t*), the loss is calculated across all possible corruptions of the tail entity, effectively capturing the model’s ability to discriminate between true and false triples. The choice of CEL over the conventional Negative Sampling (NS) approach is supported by recent comprehensive studies in knowledge graph embedding literature. Notably, Kamigaito et al.(65) demonstrated that CEL achieves stronger model fitting to training data compared to NS, attributing this to CEL’s inherent divergence and convexity properties. This observation is further corroborated by Ali et al’s (51) extensive experimentation in the largest scale benchmark study of Knowledge Graph Embedding Models (KGEMs) to date, comprising 28,000 GPU hours of testing—which revealed CEL’s consistently high performance compared to high variance in NS and Binary Cross-Entropy (BCE). The empirical evidence for CEL’s effectiveness and its theoretical advantages motivates our choice.

### Hyperparameter Optimization

Within all three training phases, Optuna(66) is employed for systematic hyperparameter optimization (HPO). Optuna’s optimization process utilizes Tree-structured Parzen Estimators (TPE) sampler for efficient parameter space navigation.

Table 2 illustrates all knowledge graph embedding mehtods hyper-parameter space, including learning rate spanning three orders of magnitude (10^−5^ to 10^−3^), varied batch sizes (512 to 4096), and embedding dimensions (256 to 2048). For the finetuning, we maintain the same search space except for embedding dimension, which remains fixed to match the pretrained model architecture.

**Table 2.** KGEM Hyperparameter Search Space.

| Hyperparameter | Range | Step |
| --- | --- | --- |
| Learning Rate | $[1 \times 10^{-5}, 1 \times 10^{-3}]$ | Log |
| Batch Size | {512, 1024, 2048, 4096} | Discrete |
| $\lambda_{n3}$ | [0.1, 0.5] | Uniform |
| Embedding Dim. | {256, 512, 1024, 2048} | Discrete |
| Epochs | {10, 20, 30, 40, 50, 60, 70} | Discrete |
| Early Stopping | Constant | Patience = 3 |

Table 3 presents the classifier hyperparameter space that is carefully selected to accommodate each classifier’s unique characteristics. For tree-based classifiers (Decision Tree, Random Forest, XGBoost), we focus on structural parameters like tree depth and number of estimators. Support Vector Machines explore various kernel functions with corresponding hyperparameters. Neural network configurations (MLP) consider different architectures through hidden layer sizes and activation functions.

**Table 3.** Classifier Hyperparameter Search Spaces.

| Model | Parameter | Range |
| --- | --- | --- |
| Decision |  |  |
| Tree | max depth | [1, 50] |
|  | min samples split | [2, 20] |
|  | min samples leaf | [1, 20] |
| Random |  |  |
| Forest | estimators | [40, 500] |
|  | max depth | [1, 25] |
|  | min split | [2, 20] |
|  | criterion | gini, entropy |
| SVM | kernel | linear, poly, rbf |
| | C, gamma | $[10^{-2}, 10^3]$, $[10^{-5}, 10^5]$ |
| KNN | neighbors | [1, 30] |
|  | weights, p | {uniform, distance}, [1, 2] |
| Logistic |  |  |
| Regression | C | $[10^{-5}, 10^2]$ |
|  | solver | newton-cg, lbfgs, liblinear |
|  | penalty | l1, l2, elasticnet |
| XGBoost | estimators | [50, 500] |
|  | max depth | [3, 20] |
| | learning rate | $[10^{-4}, 0.3]$ |
| MLP | hidden layers | (50), (100), (50,50) |
|  | activation | relu, tanh, logistic |
|  | solver | adam, sgd, lbfgs |
| | learning rate | $[10^{-4}, 0.1]$ |

## Results

This section presents a detailed analysis of the framework’s performance, which has been systematically gauged through three distinct and progressive training phases. Each phase is evaluated to assess different aspects of the framework’s capabilities and robustness. In addition, it presents a detailed case study that is done to determine BIND’s potential to predict unknown interactions between drugs and phenotypes.

### Knowledge Graph Embedding Methods Performance Analysis

Table 4’s pretrained column presents 11 distinct knowledge graph embedding methods performance on 30 distinct interaction types. These models performance for Hits@1,3,5,10, and 50 is given here. For the first phase, where the methods are trained with all 30 interactions, evaluation is performed on each relation separately. The primary objectives of this evaluation are to analyze performance across interaction types and to shortlist the top methods for the next phase.

**Table 4.** Performance comparison of 30 different relations using 16 models in terms of F1-score.

|  | Pretrained |  |  |  |  |  |  |  |  |  |  | Finetuned |  |  |  |  |
| --- | --- | --- | --- | --- | --- | --- | --- | --- | --- | --- | --- | --- | --- | --- | --- | --- |
|  | HAKE | DistMult | DualE | HolE | PairRE | QuatE | Simple | TransE | Complex | TripleRE | RotatE | DistMult | TripleRE | PairRE | HolE | DualE |
| Protein-Protein | 0.0209 | 0.0315 | 0.0287 | 0.0386 | 0.0347 | 0.0295 | 0.0318 | 0.0271 | 0.0295 | 0.0338 | 0.0298 | 0.0418 | 0.0429 | 0.0421 | 0.0416 | 0.0424 |
| Disease-Phenotype(Positive) | 0.0229 | 0.0595 | 0.0552 | 0.0657 | 0.0624 | 0.0530 | 0.0539 | 0.0345 | 0.0534 | 0.0634 | 0.0510 | 0.0706 | 0.0716 | 0.0709 | 0.0697 | 0.0707 |
| Bioprocess-Protein | 0.0394 | 0.1180 | 0.1120 | 0.1310 | 0.1240 | 0.1090 | 0.1080 | 0.0567 | 0.1100 | 0.1270 | 0.1100 | 0.1400 | 0.1390 | 0.1370 | 0.1310 | 0.1290 |
| Cellcomp-Protein | 0.1110 | 0.1610 | 0.1450 | 0.1610 | 0.1610 | 0.1490 | 0.1540 | 0.1060 | 0.1490 | 0.1620 | 0.1290 | 0.1780 | 0.1730 | 0.1770 | 0.1560 | 0.1460 |
| Drug-Effect | 0.0118 | 0.0310 | 0.0229 | 0.0282 | 0.0333 | 0.0274 | 0.0320 | 0.0182 | 0.0292 | 0.0298 | 0.0281 | 0.0310 | 0.0297 | 0.0332 | 0.0313 | 0.0291 |
| Disease-Disease | 0.0731 | 0.2560 | 0.2320 | 0.2630 | 0.2520 | 0.1820 | 0.2220 | 0.1270 | 0.2080 | 0.2520 | 0.2200 | 0.2890 | 0.2870 | 0.2820 | 0.2690 | 0.2660 |
| Phenotype-Phenotype | 0.0396 | 0.3490 | 0.3080 | 0.3380 | 0.3360 | 0.2400 | 0.3160 | 0.1400 | 0.2770 | 0.3350 | 0.2910 | 0.3740 | 0.3730 | 0.3640 | 0.3550 | 0.3470 |
| Drug-Protein | 0.0746 | 0.1920 | 0.1840 | 0.1990 | 0.1940 | 0.1760 | 0.1830 | 0.0813 | 0.1830 | 0.1880 | 0.1900 | 0.2130 | 0.2090 | 0.2100 | 0.2090 | 0.1990 |
| Anatomy-Anatomy | 0.0910 | 0.4860 | 0.4170 | 0.4610 | 0.4850 | 0.2400 | 0.4140 | 0.1950 | 0.3550 | 0.3410 | 0.3370 | 0.5020 | 0.5000 | 0.5000 | 0.4910 | 0.4890 |
| Anatomy-Protein(Absent) | 0.0882 | 0.1860 | 0.1740 | 0.1820 | 0.1870 | 0.1720 | 0.1790 | 0.0930 | 0.1750 | 0.1930 | 0.1710 | 0.2240 | 0.2210 | 0.2170 | 0.1940 | 0.1800 |
| Bioprocess-Bioprocess | 0.0560 | 0.2660 | 0.2430 | 0.2850 | 0.2670 | 0.1930 | 0.2380 | 0.1420 | 0.2160 | 0.2710 | 0.2300 | 0.3070 | 0.3070 | 0.3060 | 0.2860 | 0.2730 |
| Pathway-Protein | 0.0703 | 0.1810 | 0.1730 | 0.1900 | 0.1830 | 0.1640 | 0.1700 | 0.1020 | 0.1690 | 0.1820 | 0.1600 | 0.1860 | 0.1820 | 0.1830 | 0.1880 | 0.1790 |
| Contraindication | 0.0221 | 0.0654 | 0.0569 | 0.0692 | 0.0679 | 0.0584 | 0.0611 | 0.0299 | 0.0578 | 0.0678 | 0.0598 | 0.0652 | 0.0677 | 0.0679 | 0.0727 | 0.0666 |
| Cellcomp-Cellcomp | 0.0824 | 0.3910 | 0.3380 | 0.3590 | 0.3740 | 0.2530 | 0.3510 | 0.1560 | 0.2940 | 0.3270 | 0.3050 | 0.4190 | 0.4190 | 0.4130 | 0.4020 | 0.3910 |
| Indication | 0.0357 | 0.2030 | 0.1850 | 0.1980 | 0.1970 | 0.1760 | 0.1750 | 0.0525 | 0.1710 | 0.2050 | 0.1600 | 0.2170 | 0.2130 | 0.2080 | 0.1960 | 0.2060 |
| Molfunc-Molfunc | 0.1010 | 0.3810 | 0.3640 | 0.3800 | 0.3790 | 0.2780 | 0.3520 | 0.1600 | 0.3290 | 0.3570 | 0.3310 | 0.4050 | 0.4010 | 0.3960 | 0.3850 | 0.3770 |
| Pathway-Pathway | 0.0578 | 0.3780 | 0.3130 | 0.3550 | 0.3710 | 0.2870 | 0.3700 | 0.1570 | 0.2760 | 0.3830 | 0.2710 | 0.3970 | 0.3990 | 0.3840 | 0.3600 | 0.3850 |
| Disease-Protein | 0.0411 | 0.1090 | 0.1060 | 0.1240 | 0.1160 | 0.1060 | 0.1010 | 0.0433 | 0.1030 | 0.1170 | 0.1010 | 0.1310 | 0.1320 | 0.1390 | 0.1310 | 0.1310 |
| Anatomy-Protein-Present | 0.0221 | 0.0246 | 0.0204 | 0.0258 | 0.0241 | 0.0181 | 0.0210 | 0.0177 | 0.0200 | 0.0207 | 0.0210 | 0.0270 | 0.0273 | 0.0272 | 0.0266 | 0.0257 |
| Molfunc-Protein | 0.1300 | 0.1950 | 0.1790 | 0.1970 | 0.1970 | 0.1770 | 0.1830 | 0.1230 | 0.1800 | 0.2000 | 0.1770 | 0.2160 | 0.2160 | 0.2140 | 0.1960 | 0.1780 |
| off-label use | 0.0290 | 0.2670 | 0.1960 | 0.2380 | 0.2560 | 0.1930 | 0.2000 | 0.0352 | 0.1860 | 0.2580 | 0.1990 | 0.2910 | 0.2960 | 0.2980 | 0.2580 | 0.2680 |
| disease_phenotype_negative | 0.0518 | 0.3341 | 0.1723 | 0.2880 | 0.3300 | 0.1497 | 0.1970 | 0.0686 | 0.1897 | 0.3006 | 0.1880 | 0.4726 | 0.4446 | 0.4604 | 0.4000 | 0.4045 |
| phenotype_protein | 0.0696 | 0.2418 | 0.2045 | 0.2300 | 0.2270 | 0.1907 | 0.2002 | 0.0709 | 0.1974 | 0.2374 | 0.2015 | 0.3059 | 0.3028 | 0.3043 | 0.2735 | 0.2705 |
| exposure_protein | 0.0620 | 0.2809 | 0.2037 | 0.2060 | 0.2340 | 0.1777 | 0.2147 | 0.0827 | 0.1944 | 0.2406 | 0.2202 | 0.2991 | 0.2886 | 0.2923 | 0.2923 | 0.2599 |
| exposure_disease | 0.0372 | 0.1839 | 0.1319 | 0.1480 | 0.1660 | 0.1177 | 0.1383 | 0.0560 | 0.1348 | 0.1699 | 0.1333 | 0.2017 | 0.1893 | 0.1859 | 0.2666 | 0.1539 |
| exposure_exposure | 0.0902 | 0.2033 | 0.1376 | 0.1610 | 0.1810 | 0.1271 | 0.1650 | 0.1067 | 0.1434 | 0.1715 | 0.1555 | 0.2047 | 0.1882 | 0.1916 | 0.1916 | 0.1555 |
| exposure_bioprocess | 0.0595 | 0.2360 | 0.1668 | 0.1610 | 0.2120 | 0.1457 | 0.1790 | 0.0742 | 0.1725 | 0.2077 | 0.1914 | 0.2508 | 0.2290 | 0.2319 | 0.1586 | 0.1940 |
| exposure_molfunc | 0.2045 | 0.5746 | 0.2923 | 0.2550 | 0.5000 | 0.3272 | 0.3344 | 0.1293 | 0.3369 | 0.4806 | 0.3541 | 0.5850 | 0.4882 | 0.5237 | 0.3414 | 0.4620 |
| exposure_cellcomp | 0.1516 | 0.5900 | 0.4413 | 0.4360 | 0.5470 | 0.4643 | 0.6450 | 0.1549 | 0.4786 | 0.4249 | 0.4581 | 0.5900 | 0.2876 | 0.5467 | 0.2182 | 0.4410 |
| drug_drug | 0.0196 | 0.0186 | 0.0164 | 0.0198 | 0.0181 | 0.0135 | 0.0161 | 0.0095 | 0.0144 | 0.0179 | 0.0149 | 0.0196 | 0.0180 | 0.0185 | 0.0201 | 0.0181 |

The evaluation (Table 4) reveals distinct patterns in the methods’ ability to capture biomedical relationships. HolE(43), DistMult(35), PairRE(36), TripleRE(37), and DualE(39) consistently emerge as the top performers across all evaluation metrics. These methods have achieved superior performance through their ability to learn features from sparse relations effectively despite the two overbearing dominant relations(anatomy-protein(present) and drug-drug). HolE’s circular correlation operation proves particularly adept at capturing the highly bidirectional nature of PrimeKG’s relations, where nearly all relationship pairs (h,r,t) and (t,r’,h) coexist. DistMult’s strong performance can be attributed to its simple multiplicative interactions aligning well with the predominantly straightforward complexity of the most biological relations, requiring neither asymmetric nor compositional properties. PairRE and TripleRE (both extensions of TransE) excel through their simple methodology and relation-specific projections. DualE’s hybrid (rotation and translation both) quaternion-based approach seems to successfully balance modeling of both symmetric (such Drug-Drug) and hierarchical (such Anatomy-Protein) relationships.

TransE(34), SimplE(44), and HAKE(42) perform poorly on PrimeKG due to architectural mismatches. TransE struggles with many-to-many relationships common in biological data, while HAKE’s hierarchical design may have faltered on the prevalent non-hierarchical Drug-Drug interactions. QuatE(38), ComplEx(40), and RotatE(41) show good but suboptimal performance, as their sophisticated mathematical approaches prove unnecessary for PrimeKG’s relations. Successful learning in large scale biomedical knowledge graphs appears to depend more on handling relation imbalance, bidirectionality, and varying cardinalities rather than complex geometric transformations.

The results reveal significant variation in model performance across different relation types. As shown in Table 4, the most frequent relations in PrimeKG exhibit some of the lowest MRR scores. This suggests that the densely connected protein-protein interactions and complex many-to-many interactions in anatomy-protein(present) are the most difficult to effectively learn, presenting a significant challenge for scalable embedding methods. The highest MRR scores are achieved on more specialized relations such as Anatomy-Anatomy (0.486), Pathway-Pathway (0.383), and Cellular Component-Cellular Component (0.391). This pattern persists across all top-performing models, with HolE, PairRE, TripleRE and DistMult showing particularly strong performance on these specialized relations. The superior performance on sparse relations might be attributed to their more focused semantic scope and potentially simpler underlying patterns. For instance, anatomy-anatomy interaction follows a clear hierarchical structures, while protein localization patterns (Anatomy-Protein) involve complex many-to-many mappings that prove challenging to model effectively. These findings underscore the need for generalizable and simple architectural innovations that can better handle both high-frequency relations with complex interaction patterns and sparse relations with simpler underlying structures.

### Impact of Relation-Specific Fine-Tuning on Knowledge Graph Embeddings Performance

This section presents an analysis of the extent to which relation-specific fine-tuning improves the performance of pre-trained models. This phase only utilizes the 5 top-performing methods namely HolE(43), DistMult(35), PairRE(36), TripleRE(37), and DualE(39).

The top 5 methods performance across the 30 interaction types in terms of Hits@1, 3, 5, 10, and 50 is available here. Table 4’s finetuned column illustrates the MRR performance difference between the 5 finetuned methods and the 10 pretrained methods from the previous phase. A high level analysis of Table 4 reveals that most relatively substantial improvements are observed in structurally complex (many-to-many) interactions. Protein-Protein interactions show relative improvements of 21.3% (from 0.0347 to 0.0421 with PairRE) and 26.9% (from 0.0338 to 0.0429 with TripleRE). Similarly, Disease-Disease relationships see notable improvements across all models, with DistMult showing the largest gain (12.9% improvement, from 0.256 to 0.289). These improvements in complex biological relationships indicate that finetuning helps models better capture domain-specific interaction patterns. Despite the overall increase in model performance, some persistent challenges remain. While we see improvements with focused training, high frequency relations like Anatomy-Protein (Present) still exhibit a low MRR performance (highest achieved is 0.272). This pattern suggests that effectively modeling these types of relationships requires more than just focused training, and may need fundamental architectural improvements to the model.

The relative performance ranking of models remains largely stable after finetuning, with PairRE, TripleRE, and DistMult consistently achieving the best results across most relations. However, the gap between models narrows after finetuning, particularly for specialized biological relationships. This convergence suggests that relation-specific optimization helps compensate for architectural differences between models, though the underlying strengths and weaknesses of each approach remain evident in the final performance.

### Interaction Prediction Performance

Knowledge graph embedding methods have demonstrated acceptable performance across most relation types, but there remains significant room for predicting more accurately. Combining the embedding spaces produced by these methods with machine learning classifiers may enable better identification of previously unknown interactions. To develop effective predictive pipelines, it is essential to analyze which combinations of embedding methods and machine learning classifiers produce optimal performance. To perform this analysis, we conducted extensive experimentation. The detailed performance metrics including precision, recall, and accuracy for the top performing methods for each relation are provided in Supplementary File 1, whereas the top-performing predictive pipeline for each relation type is presented in Figure 3. The interaction prediction results highlight the effectiveness of our two-stage embedding approach, which achieved consistently high F1-scores across diverse biological relationships.

**Fig. 3.**
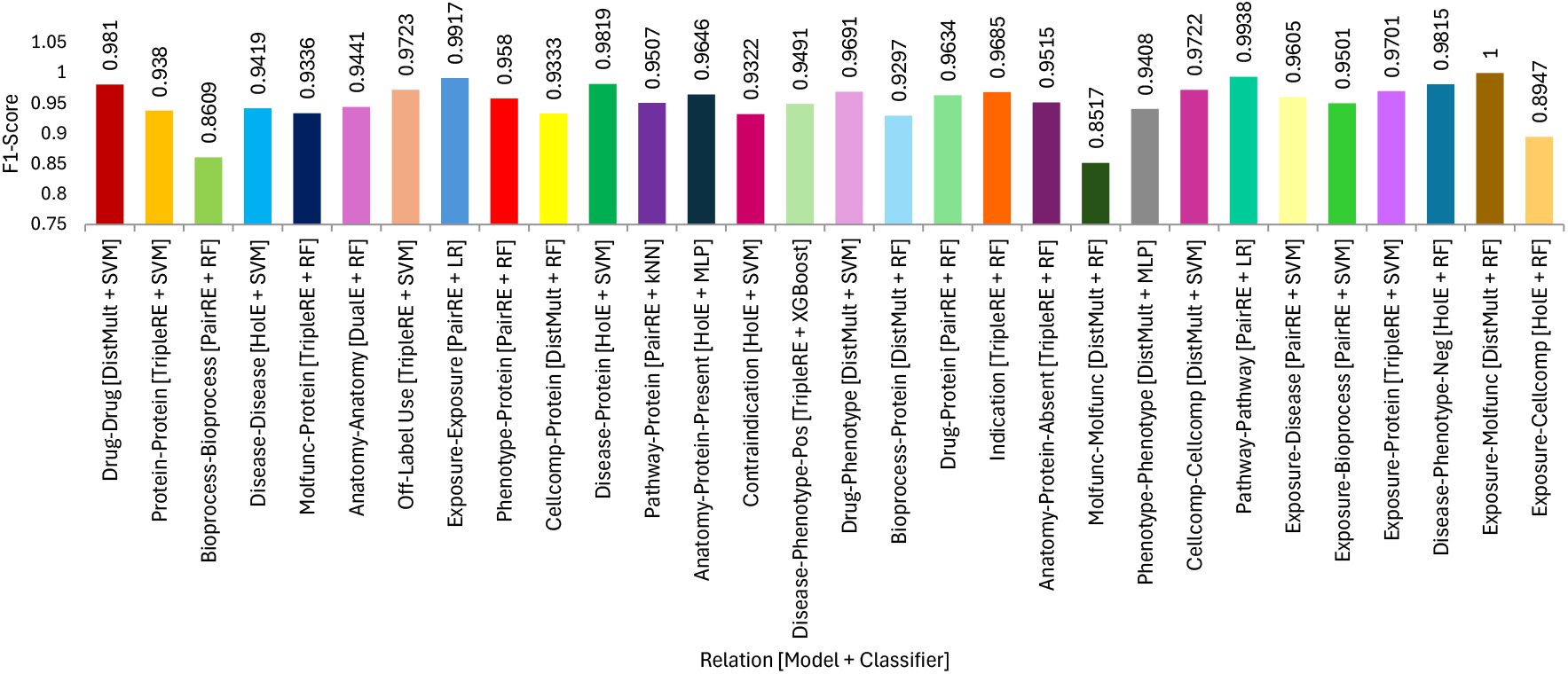
Best Embedding Method + Classifier Combination For Each Relation

Particularly strong performance is observed for drug-related predictions, with Drug-Drug interactions achieving an F1-score of 0.9810 using DistMult embeddings with an SVM classifier, and Drug-Effect predictions reaching 0.9940 with PairRE embeddings and SVM. These results are especially significant given the high volume of drug-related data in PrimeKG and the critical importance of accurate drug interaction predictions in clinical applications.

Complex biological relationships, such as Protein-Protein interactions and Disease-Protein associations, show robust performance (F1-scores of 0.9380 and 0.9819 respectively), indicating that the finetuned embeddings effectively capture the intricate patterns in these interactions. The high-frequency Anatomy-Protein-Present interactions achieves a strong F1-score of 0.9646 using HolE embeddings with an MLP classifier, despite its challenging many-to-many nature that posed difficulties in the embedding phase. The results also reveal that smaller, specialized relation types benefit from different classifier architectures. For instance, Pathway-Protein predictions achieve optimal performance (F1=0.9507) using k-Nearest Neighbors with PairRE embeddings, suggesting that local neighborhood structure is particularly informative for pathway-related predictions.

These results validate that our predictive pipelines, which combine relation-specific embedding fine-tuning with tailored classifier selection, are effective for biological interaction prediction. The consistently high F1-scores across diverse relation types demonstrate their ability to capture varied semantic patterns in PrimeKG(27).

### BIND: Biological Interaction Network Discovery Web Application

Building on the success of our framework in identifying highly accurate predictive pipelines (Figure 3), we introduce the BIND web application (https://sds-genetic-interaction-analysis.opendfki.de/). BIND serves as a robust platform for the bioinformatics community, offering real-time exploration of high-confidence predictions across diverse biological relationships.

To ensure practical application and broaden the biological interaction landscape, we consolidate the optimal predictive pipeline for each relation type into a unified platform. This enables researchers to seamlessly investigate interactions at scale. Our pipelines were employed to evaluate 2.34 billion potential novel interaction pairs (2,338,684,118 combinations) across all interaction types. Predictions with confidence scores below 70% were excluded, and the remaining high-confidence interactions were integrated into the web application, facilitating real-time querying and analysis of these potentially novel interactions. To evaluate the practical utility of BIND, we conducted a case study on drug-phenotype interactions.

### Drug-Phenotype Interaction Prediction: A Case Study

Drug-phenotype interactions are a vital subset of biomedical relationships with significant clinical implications. This case study focuses on this relationship due to its potential for advancing drug repurposing and adverse effect prediction. Moreover, the drug-effect relation exhibited low performance during the initial training phases (Table 4), making it an ideal conservative test case—success here would strongly validate our approach without overstating the framework’s capabilities.

Our approach successfully identified well-documented drug-phenotype associations. Table 5 highlights high-confidence predictions consistent with established pharmacological mechanisms (e.g., Propranolol-Wheezing, confidence 0.9998, attributed to beta-blockade effects) but absent from the training data. This demonstrates the model’s ability to rediscover known interactions, indicating that other high-confidence predictions may uncover novel, biologically plausible associations.

**Table 5.** Predicted Drug-Phenotype Associations with Strong Literature Support.

| Drug Class | Drug | Phenotype | Confidence | Literature Support |
| --- | --- | --- | --- | --- |
| Beta Blocker | Propranolol | Wheezing | 0.9998 | Well-documented beta-blockade induced bronchospasm (67) |
| 5-alpha-reductase Inhibitor | Dutasteride | Abnormal spermatogenesis | 0.9997 | Established DHT inhibition effects on male reproduction (68) |
| Anticholinergic | Methscopolamine | Nephropathy | 0.9998 | Known effects on renal function via muscarinic blockade (69) |
| Antiretroviral | Efavirenz | Hepatic necrosis | 0.9927 | Well-documented hepatotoxicity in HIV treatment (70) |
| Chemotherapy | Dacarbazine | Chronic leukemia | 0.9626 | Known hematological effects of alkylating agents (71) |
| Anticonvulsant | Zonisamide | Epistaxis | 0.9899 | Documented coagulation effects (72) |
| Antipsychotic | Flunitrazepam | Hypothyroidism | 0.9897 | Established endocrine system interactions (73) |
| Antiemetic | Granisetron | Encephalopathy | 0.9896 | Known 5-HT3 effects on CNS (74) |
| Antineoplastic | Fludarabine | Immunodeficiency | 0.9531 | Well-documented immunosuppressive effects (75) |
| NSAID | Ketorolac | Limitation of neck motion | 0.9902 | Known musculoskeletal effects (76) |
| Corticosteroid | Fluocinonide | Macular hole | 0.9795 | Documented ocular complications (77) |
| Antihistamine | Hydroxyzine | Reticulocytosis | 0.9904 | Known hematological effects (78) |
| Opioid Antagonist | Dextromethorphan | Myoclonus | 0.9929 | Established neurological side effects (79) |
| Muscle Relaxant | Baclofen | Respiratory depression | 0.9912 | Known effect on respiratory centers (80) |
| Antifungal | Griseofulvin | Photosensitivity | 0.9901 | Documented phototoxic reactions (81) |
| Antimalarial | Chloroquine | Retinopathy | 0.9899 | Well-established ocular toxicity (82) |
| Antiplatelet | Clopidogrel | Bleeding time increased | 0.9898 | Known effect on coagulation (83) |
| Bronchodilator | Albuterol | Tachycardia | 0.9897 | Documented beta-2 cardiac effects (84) |
| Antidepressant | Escitalopram | Abnormal heart rhythm | 0.9445 | Known cardiac conduction effects (85) |
| Growth Hormone | Octreotide | Decreased liver function | 0.9411 | Established hepatic metabolic effects (86) |

Analysis of confidence score distributions revealed a clear bimodal pattern: a sharp peak above 0.99 for many known predictions. This separation suggests the model effectively model likely pharmacological effects. The prevalence of literature-supported predictions among high-confidence results, validates our methodological approach and the potential of embedding-based methods for extending drug-phenotype knowledge. Wet lab experimental validation of these unknown predictions could expand our understanding of drug effects and reveal new therapeutic opportunities.

## Conclusion

This research investigates the efficacy of Knowledge Graph Embedding Methods (KGEMs) in generating high-quality embeddings for biological entities. Our findings demonstrate that a two-stage training approach, encompassing initial training on diverse relational data followed by fine-tuning on specific relation types, significantly enhances the quality of the generated embeddings. Furthermore, extensive experimentation across 30 distinct relation types involving 11 KGEMs revealed that simpler model architectures often effectively capture the inherent characteristics of biological interactions, surpassing the performance of more complex models. While KGEMs exhibit strong capabilities in predicting interactions between various biological entities, their integration with machine learning classifiers significantly enhances predictive performance. Although a universal predictive pipeline achieving optimal performance across all interaction types remains elusive, our research successfully identified specific combinations of KGEMs and classifiers for each interaction type, resulting in F1 scores ranging from 90% to 99%. To facilitate the practical application of these findings, the BIND-web application was developed, enabling the prediction of unknown interactions between 10 distinct entity types across 30 different relation types. A comprehensive evaluation of the BIND-web application with a Drug-Phenotype Interaction Prediction case study demonstrated its significant potential for accurate prediction of novel interactions. A promising future direction for this research lies in extending the scope of the web application to identify potential interaction pathways connecting diverse biological entities.

